# MXRA7 Alleviates Epididymitis from Exercise-Induced Fatigue by Inhibiting Pyroptosis

**DOI:** 10.64898/2026.02.11.705342

**Authors:** Kun-Yang Tang, Xiao-Cui Jiang, Zhi-Peng Fang, Xin Hu, Jia-Sen Liu, Xiao-Ming Yu, Min Zhao, Ying Liu, Ji-Gang Cao, Yan-Yan Zhou, Min Xiao

## Abstract

**Aims:** To explore exercise-induced fatigue (EIF)’s effects on the male reproductive system and MXRA7’s regulatory role herein.

**Methods:** We recruited EIF volunteers for semen/serum tests, established a mouse EIF model via weight-loaded swimming to assess epididymal segmental injury, and constructed pyroptosis models of PC-1/DC-2 cells. Public database transcriptomic analysis identified MXRA7 expression and enriched pathways in epididymitis; MXRA7’s function was verified via its knockdown/overexpression in DC-2 cells. PKCα-MXRA7 association was explored by phosphorylation assays and CO-IP, and sperm incubation experiments evaluated MXRA7’s effect on sperm function.

**Results:** EIF impaired human sperm motility, reduced mouse sperm quality and induced epididymitis with segment-specific pyroptosis. MXRA7 expression differed in PC-1/DC-2 cells and correlated with pyroptosis; it was phosphorylated by PKCα, inhibited the NF-κB pathway to alleviate inflammation, and mitigated pyroptosis-induced sperm motility damage.

**Conclusion:** EIF induces epididymal epithelial pyroptosis and epididymitis, and MXRA7 exerts a protective effect mainly in caudal epididymal cells by alleviating pyroptosis, thus reducing sperm quality damage.

## 1 Introduction

Exercise-induced fatigue (EIF) refers to a phenomenon characterized by the decline of physical functions caused by excessive and inappropriate exercise(Ament and Verkerke, 2009). Transient and mild EIF usually resolves spontaneously without special intervention, which can be regarded as a normal physiological mechanism of the body and is extremely common. However, long-term and severe EIF can impair the body’s health status and even induce organic diseases(Tang et al., 2025), among which the male reproductive system is also affected. Current evidence, including the findings from our preliminary studies, has indicated the effect of EIF on the male reproductive system, particularly its association with oligoasthenospermia(Tang et al., 2025, Roberts et al., 1993, Cupka and Sedliak, 2023, Jóźków and Rossato, 2017), but research in this field remains insufficient.

Epididymitis is a common intrascrotal inflammatory condition, with an incidence of approximately 400 cases per 100,000 males worldwide annually(Fijak et al., 2018). It is frequently caused by ascending bacterial infection of the genitourinary tract, such as sexually transmitted diseases (STDs) or urinary tract infections (UTIs) induced by multiple factors. Currently, persistent oligoasthenospermia and azoospermia occur in up to 40% of males, and epididymitis is recognized as one of the primary etiological factors(Da Silva et al., 2025). Considering the unique systemic inflammatory response induced by EIF, the physical friction exerted on reproductive appendages (including the epididymis) during exercise(Jóźków and Rossato, 2017), and the potential immunomodulatory effects of EIF, as well as the susceptibility of the epididymis to systemic stress and strain(Moon et al., 2024), we hypothesized that the oligoasthenospermia phenotype induced by long-term EIF may be associated with the development of epididymitis, which formed the rationale for the present study.

Pyroptosis is an inflammatory form of programmed cell death(Burdette et al., 2021). Lipopolysaccharide (LPS) secreted by bacteria (e.g., Gram-negative bacteria) and endogenous adenosine triphosphate (ATP) released by host cells upon bacterial infection form a synergistic inflammatory signal, which induces mitochondrial dysfunction and simultaneously activates gasdermin D (GSDMD) to form annular pores in the plasma membrane, ultimately leading to cell death(Deo et al., 2020, Vasudevan et al., 2023). The pathogenic process of epididymitis shares similarities with the pyroptosis mechanism; therefore, pyroptosis may also be involved in EIF-induced epididymitis. Pyroptosis of epithelial cells that are in direct contact with sperm is likely to trigger inflammatory changes in the intraluminal microenvironment of the epididymis, thereby exerting adverse effects on sperm.

Although the epithelial cells lining the caput, corpus, and cauda epididymidis share similar components, they exhibit distinct differences in cellular morphology and function. In epididymitis, the caput epididymidis typically displays a higher degree of inflammatory activity than the cauda epididymidis(Da Silva et al., 2025), and a similar pattern has been observed in EIF-induced epididymitis. While previous studies have proposed that this phenomenon is attributed to the structural similarity between the caput epididymidis and the seminiferous tubules of the testis, as well as the more complex physical architecture of the caput region, the specific underlying mechanisms remain poorly elucidated. Considering the functional heterogeneity of epithelial cells across different epididymal segments, it is plausible that certain regulatory factors may modulate the inflammatory activity of cells in distinct epididymal regions.

Matrix remodeling associated 7 (MXRA7) is a secreted functional protein, and previous studies have demonstrated that it also acts as an autocrine factor(Shen et al., 2023). MXRA7 has been shown to be associated with inflammatory signaling pathways such as NF-κB(Sun et al., 2023). In our study, we found that MXRA7 expression exhibits segment-specific differences across the caput, corpus, and cauda epididymidis, which is positively correlated with the degree of inflammation. Therefore, we hypothesized that MXRA7 may have a certain association with epididymitis. Despite the paucity of existing research on MXRA7, scattered studies have revealed that it exerts prominent effects in inflammatory and immune responses (Sun et al., 2024), and it is also implicated in cell proliferation, division, differentiation, and wound repair(Shen et al., 2023). Based on these findings, we proposed that MXRA7 may exert a protective role in epididymitis, which formed the basis of the present study.

## 2 Materials and Methods

### 2.1 Reagents

Vitamin C, lipopolysaccharide (LPS), phorbol 12-myristate 13-acetate (PMA), BIM 1 (Orileaf, China); Tunel, IL-6, IL-1β, and TNF-α assay kits; 8-OhdG; FITC-labeled Arachis hypogaea agglutinin; Anti-MXRA7 antibody, Anti-p-PKCα (Ser657/Tyr658) Antibody (Sigma-Aldrich); Anti-Hexokinasel antibody; Anti-COX IV antibody; Anti-α-Tubulin antibody; Anti-NLRP3 antibody, Anti-Caspase-1 antibody, Anti-GSDMD antibody, Anti-p-NF-κB p65 antibody, Anti-NF-κB p65 antibody, Anti-p-NF-κB IκBα antibody, Anti-NF-κB IκBα antibody, Anti-NF-κB p52/p100 antibody , Anti-PKCα antibody, Anti-pan-PKC antibody , HRP-conjugated Goat Anti-Rabbit IgG(H+L), (Wuhan Sanying, China); Phos-tag SDS-PAGE (FUJIFILM Wako, Japan); DAPI staining reagent (Wuhan Servicebio, China).

### 2.2 Human Subjects

Human semen and serum samples were collected from volunteers aged 25–40 years at the Guanggu Campus of Hubei Provincial Hospital of Traditional Chinese Medicine. All volunteers attended the hospital either voluntarily or upon invitation for reproductive medicine diagnosis or treatment, and were all native speakers of simplified Chinese. Sample evaluation was performed in accordance with the principles outlined in the 6th edition of the World Health Organization (WHO) Laboratory Manual for the Examination and Processing of Human Semen (World Health Organization, 2021). All experiments were conducted in compliance with the Declaration of Helsinki of the World Medical Association. The study was approved by the Institutional Ethics Committee (approval No.: HBZY2023-C10-02), and written informed consent was obtained from each participant.

We initially collected sperm data from 169 male participants, and 45 volunteers remained after the implementation of inclusion and exclusion criteria, including 15 patients with exercise-induced fatigue (EIF) who consented to both sperm and serum sample collection, and 30 healthy volunteers, among whom 15 agreed to serum sample donation.The inclusion criteria for EIF patients were defined as follows: 1)Male subjects aged 18–45 years who visited the hospital voluntarily or upon invitation for sperm quality testing due to reproductive health consultation and fertility preparation needs, with signed informed consent and willingness to cooperate with the study procedures. 2)A confirmed history of EIF exposure within 1 week prior to testing: for regular exercisers, the intensity of a single exercise session was increased by more than 50% relative to their routine level for 3 consecutive days; for irregular exercisers, sudden high-intensity exercise was performed for more than 3 consecutive days. 3)A score of ≥ 6 on the Fatigue Severity Scale (FS-14) and a score of ≥ 7 on the Borg Rating of Perceived Exertion (RPE) scale immediately after exercise; persistent fatigue symptoms were reported daily within the last 2 weeks, which were exacerbated after exercise, could not be completely relieved even after more than 30 minutes of rest, and persisted until the time of testing. 4)Provision of a complete semen sample with a volume of ≥ 1.5 mL, without missing baseline data.

The inclusion criteria for healthy volunteers were as follows: 1)Male subjects aged 18–45 years who underwent sperm quality testing at the hospital during the same period, with an age difference of ± 5 years relative to the EIF patient group, and who provided informed consent and voluntarily agreed to participate in the study. 2)No history of EIF exposure within the preceding two weeks; a score of < 3 on the Fatigue Severity Scale (FS-14) and a score of < 4 on the Borg Rating of Perceived Exertion (RPE) scale immediately after exercise. 3)Normal sperm parameters and no history of definite diseases, including no medical history of reproductive system disorders, endocrine system disorders, or chronic systemic diseases. 4)Compliance with the same semen sample collection standards as the EIF patient group, with complete and traceable baseline data.

Exclusion criteria were as follows: 1)A history of organic diseases or surgical interventions related to the reproductive system. 2)Presence of systemic diseases, including endocrine/metabolic disorders (e.g., diabetes mellitus, thyroid dysfunction), autoimmune diseases, chronic infectious diseases (e.g., tuberculosis, human immunodeficiency virus (HIV) infection), and hepatic or renal insufficiency. 3)Exposure to interfering medications or lifestyle factors: use of drugs affecting sperm quality (e.g., antidepressants, hormonal agents) within the preceding 3 months; smoking ≥ 10 cigarettes per day for ≥ 1 consecutive year; alcohol dependence or substance abuse; or administration of hormone supplements and high-dose antioxidants. 4)Presence of confounding factors for fatigue diagnosis: fatigue caused by non-exercise related etiologies (e.g., anemia, sleep duration < 6 hours per day for ≥ 1 consecutive month); confirmed diagnosis of depression/anxiety with ongoing medication; or incomplete or logically inconsistent completion of study scales. 5)Other confounding factors: chromosomal abnormalities, non-standardized semen sample collection, or missing baseline data.

### 2.3 Collection of Human Samples

Semen samples were obtained via masturbation following 3–5 days of sexual abstinence. After liquefaction at 37 °C for 30 min, aliquots of the semen were taken for subsequent analysis. The remaining liquefied semen was centrifuged at 400 × g for 5 min at room temperature to separate spermatozoa from seminal plasma (Barrachina et al., 2022). The supernatant seminal plasma was transferred and subjected to three consecutive rounds of centrifugation at 700 × g for 5 min at 4 °C; the resulting precipitates were discarded, and the purified seminal plasma was stored at −80 °C. The sperm pellets were resuspended in phosphate-buffered saline (PBS) and washed three times, followed by fixation in 4% PFA at room temperature for 5 min. The fixed spermatozoa were then washed again by centrifugation at 700 × g for 5 min in PBS, spread onto air-dried glass slides, and stored at −20 °C.

Serum samples were collected from fasting subjects (8–12 h of food and water deprivation) on the same day as semen sample collection. After disinfecting the venipuncture site, 3–5 mL of venous blood was drawn and allowed to clot at 25 °C for 30 min. The clotted blood was centrifuged at 3000 × g for 10 min at 4 °C to separate serum. The upper serum layer was aspirated and immediately stored at −80 °C. All procedures were completed within 2 h after blood collection.

### 2.4 Transcriptomic Analysis (RNA-seq)

Transcriptome data, including 3 samples from GSE145443 and 6 samples from GSE199903, were downloaded from the NCBI Gene Expression Omnibus (GEO) public database (https://www.ncbi.nlm.nih.gov/geo/info/datasets.html) for transcriptomic analysis. First, the expression profiles were imported and processed using the Seurat R package. Cells were filtered based on three metrics per cell: the total number of unique molecular identifiers (UMIs), the number of expressed genes, the percentage of mitochondrial gene reads, and the percentage of ribosomal gene reads. Outliers were defined as values deviating by more than three median absolute deviations (MADs) from the median value. In general, cells with excessively high total UMI counts or numbers of expressed genes are considered doublets, whereas cells with excessively high percentages of mitochondrial or ribosomal gene reads are regarded as low-quality cells that are either undergoing apoptosis or have degraded into cellular debris.

The NormalizeData function was used to standardize the raw data, followed by calculation of cell cycle scores via the CellCycleScoring function, identification of highly variable genes using FindVariableFeatures, and normalization of the data with ScaleData—this step also eliminated confounding effects of mitochondrial genes, ribosomal genes, and cell cycle phase on subsequent analyses. Linear dimensionality reduction of the expression matrix was performed using RunPCA, and the top principal components (PCs) were selected for downstream analyses. Batch effects were removed with Harmony, an algorithm that iteratively clusters similar cells from different batches in the PCA space while preserving batch diversity within each cluster. Non-linear dimensionality reduction was conducted via RunUMAP (Uniform Manifold Approximation and Projection, UMAP). Cell neighbors were identified using FindNeighbors, and cells were partitioned into distinct clusters with FindClusters. Cell annotation was performed by identifying cell types and their corresponding marker genes present in the target tissue; this was primarily achieved by querying the CellMarker database and reviewing relevant literature, with automated annotation using the SingleR software as a supplementary approach.

For RNA-seq data analysis, lowly expressed genes were filtered out with the criterion of retaining only those genes with a count > 1 in ≥ 10% of the samples. The raw count data were then normalized using the trimmed mean of M-values (TMM) method via the edgeR package and subsequently transformed into log2-counts per million (log2-CPM). Based on the median expression level of MXRA7 (Entrez ID: 439921), the samples were stratified into a MXRA7 high-expression group and a MXRA7 low-expression group. Differential expression analysis was performed using the limma package to calculate the log2 fold change (logFC) of each gene. The resulting genes were ranked by logFC values to generate a ranked gene list for gene set enrichment analysis (GSEA). The annotated gene set of version 7.0 downloaded from the Molecular Signatures Database (MsigDB) was used as the background gene set for pathway enrichment analysis between different groups, and the significantly enriched gene sets (adjusted p-value < 0.05) were sorted according to their normalized enrichment scores (NES). GSEA was further conducted using two gene sets defined by Entrez IDs: one derived from the NF-κB pathway in the Kyoto Encyclopedia of Genes and Genomes (KEGG) database, and the other a custom gene set corresponding to the canonical and non-canonical NF-κB pathways. Significance was assessed by 1,000 permutation tests, and GSEA plots illustrating the regulatory effect of MXRA7 on the NF-κB pathway were generated using the enrichplot package.

### 2.5 Animal Models and Interventions

Specific-pathogen-free (SPF) male ICR mice (weighing 30–32 g, purchased from the Hubei Provincial Center for Experimental Animals) were subjected to a 7-day acclimatization period prior to the experiment. The mice were randomly divided into three groups: the control group (CON), the exercise-induced fatigue group (EIF), and the vitamin C intervention group (Vitamin C group), with 15 mice per group. The EIF animal model was established as described previously(Yan et al., 2022), using a 4-week consecutive weight-loaded swimming training protocol. The swimming tank used for training had a water depth of 50 cm and a water temperature of (31±2) °C. During each training session, the mice were placed on the water surface, and a glass rod was gently stirred beside them to guide the mice to swim. The training lasted for 4 weeks, with the following progressive loading regimen: no weight was applied in the first week, 2% of the body weight was used as the load in the second week, 4% in the third week, and 5% in the fourth week. Each training session was terminated when the mice reached exhaustion, which was defined as the mouse failing to surface within 3 s after sinking. After the 4-week weight-loaded swimming training, all mice underwent a single session of unloaded exhaustive swimming, and the exhaustive swimming time was recorded for each mouse. Except for the CON group, mice in the other two groups received daily weight-loaded swimming training. During the 3rd–4th weeks of weight-loaded swimming training, mice in the Vitamin C group were administered vitamin C via gavage at a daily dose of 40 mg/kg body weight(Chen et al., 2022). The body weight of each mouse was recorded daily throughout the modeling period. Following the final unloaded exhaustive swimming session, the mice were retrieved, dried thoroughly, anesthetized, and subjected to blood collection. The collected blood samples were allowed to stand for 1 h, then centrifuged at 3,000 r/min for 10 min at 4 °C. The serum was separated and stored at −80 °C. The left gastrocnemius muscle and epididymides from both sides were harvested and preserved. The weight of the epididymides was measured immediately. A portion of the epididymides was punctured, placed in pre-warmed PBS at 37 °C, and incubated in a 37 °C water bath for subsequent detection. Seminal plasma and spermatozoa were separated, and sperm smears were prepared as previously described(Sacks et al., 2018). All experimental procedures were strictly performed in accordance with the Guidelines for the Care and Use of Laboratory Animals and were approved by the Institutional Animal Care and Use Committee (IACUC) of Hubei University of Chinese Medicine (Approval No.: HUCMS55714520).

### 2.6 Determination of Testosterone, BUN, LA, LDH, MDA, SOD, IL-1β, IL-6 and TNF-α

Liquid samples derived from humans or mice were assayed using enzyme-linked immunosorbent assay (ELISA) kits specific to the corresponding species, targeting testosterone, blood urea nitrogen (BUN), lactic acid (LA), lactate dehydrogenase (LDH), malondialdehyde (MDA), superoxide dismutase (SOD), interleukin-1β (IL-1β), interleukin-6 (IL-6), and tumor necrosis factor-α (TNF-α). All procedures were performed strictly in accordance with the manufacturers’ instructions.。

### 2.7 Analysis of Sperm Parameters Using an Automatic Sperm Analyzer

After complete liquefaction of human or murine semen samples, 5 μL of semen was pipetted onto a pre-warmed specialized glass slide. Sperm parameters were analyzed using the BEION Sperm Quality Analysis and Management System at 37 °C. The system captured six randomly selected microscopic fields (25 frames per second; 25 images per field), with all parameters analyzed automatically by the software. The analyzed parameters included sperm concentration (×10⁶ cells/mL), sperm motility (%), progressive motility (%), average path velocity (VAP), curvilinear velocity (VCL), straight-line velocity (VSL), amplitude of lateral head displacement (ALH), and beat-cross frequency (BCF).

### 2.8 Immunofluorescence

For cell and sperm staining: Frozen fixed sperm smears were thawed (direct exposure to ambient air was avoided during thawing to prevent contamination by condensation) or pre-prepared cell climbing slides were retrieved. The samples were permeabilized with 1% Triton X-100 and then blocked with 5% bovine serum albumin (BSA) for 1 hour. Pre-adsorbed primary antibodies were added to the slides, followed by incubation at 4 °C for 14–16 hours. After washing the slides five times with phosphate-buffered saline with Tween-20 (PBST), secondary antibodies were applied and incubated for 1 hour. Nuclei were stained with 4’,6-diamidino-2-phenylindole (DAPI) for 5 minutes, and images were visualized under a fluorescence microscope(Sacks et al., 2018). For animal tissue staining: Epididymal tissues were harvested and fixed in a universal fixative for 48 hours at room temperature, followed by paraffin embedding and sectioning. Paraffin sections were baked, dewaxed in a graded series of xylene and ethanol solutions, and then subjected to hydration and antigen retrieval. Sections were incubated with a solution containing 15% BSA and 0.5% Triton X-100 at 37 °C for 1 hour. Primary antibodies were then added at a dilution of 1:50 and incubated at 4 °C overnight. After rinsing with phosphate-buffered saline (PBS), secondary antibodies were added at a dilution of 1:1000 and incubated at 37 °C for 30 minutes. Following PBS rinses, cells were stained with specific dyes. Immediately after the final PBS rinse, sections were mounted using an anti-fluorescence quenching mounting medium, and fluorescence images were captured under a fluorescence microscope.

### 2.9 Observation of Sperm Acrosome

The processing method for semen samples used in acrosome detection was performed as described in Study(Young et al., 2024) with minor modifications. Briefly, semen samples were incubated at 37 °C for 2 hours to induce capacitation, followed by a further 1-hour incubation at 37 °C with the addition of progesterone (5 μM). The samples were then centrifuged at 700 × g for 5 minutes at room temperature, and resuspended in a hypotonic swelling medium (PBS: ddH₂O = 1:10) for 1 hour at 37 °C. After an additional washing step, the spermatozoa were resuspended and fixed in 4% PFA for 5 minutes at room temperature, then washed again by centrifugation at 700 × g for 5 minutes in PBS, and finally spread onto air-dried glass slides. For detection, the slides were incubated with 1 mg/mL FITC-labeled Arachis hypogaea agglutinin (PNA) in PBS at 4 °C for 20 minutes in the dark. The slides were rinsed with PBS, incubated with 5 mg/mL DAPI in ddH₂O for 10 seconds at room temperature, washed again with PBS, air-dried, and imaged under a fluorescence microscope.

### 2.10 Hypo-Osmotic Swelling (HOS) Test

The protocol was performed as described in Studies(Ramu and Jeyendran, 2013, Liao and O’Flaherty, 2023) with minor modifications. Processed sperm samples were gently mixed with 150 μL of hypo-osmotic swelling (HOS) solution (containing 1.5 mM fructose and 1.5 mM sodium citrate), followed by incubation at 37 °C for 30 minutes. Subsequently, 200 spermatozoa were observed under a microscope, and the number of abnormal spermatozoa (i.e., the proportion of sperm exhibiting tail swelling) was counted.

### 2.11 Histological Morphology Observation

Fixed gastrocnemius and epididymal tissues were subjected to dehydration, clearing, and wax infiltration prior to paraffin embedding. Sections with a thickness of 5 μm were cut, stained with hematoxylin and eosin (H&E), mounted with neutral resin, and observed under a light microscope.

### 2.12 Immunohistochemistry

Fixed epididymal tissues were subjected to dehydration, clearing, and wax infiltration, followed by paraffin embedding. Paraffin sections were first dewaxed and rehydrated. The staining procedure included the following steps: antigen retrieval, treatment with 3% hydrogen peroxide, demarcation of the target area, serum blocking, incubation with primary antibodies at 4 °C overnight, incubation with secondary antibodies at room temperature, color development, and counterstaining with hematoxylin. Finally, the sections were dehydrated and mounted before microscopic examination.

### 2.13 Cell Culture

The conditionally immortalized murine distal epididymal epithelial cell line (DC-2) and proximal epididymal epithelial cell line (PC-1) were purchased from Shanghai Yaji Biotechnology Co., Ltd. These cell lines were isolated from transgenic mice harboring a temperature-sensitive simian virus 40 large T tumor antigen. The culture conditions were as follows: the cells were cultured in complete Iscove’s Modified Dulbecco’s Medium (IMDM) supplemented with 10% fetal bovine serum (FBS) and 1% penicillin/streptomycin(Moon et al., 2024), in a humidified incubator at 37 °C with 5% CO₂. For pyroptosis induction(Yang et al., 2024b), the cells were seeded into 6-well plates at a density of 1×10⁶ cells/mL. After 24 hours of culture, the cells were incubated with lipopolysaccharide (LPS) at a concentration of 1 μg/mL for 5 hours, followed by treatment with adenosine triphosphate (ATP) at 5 mM for 12 hours to induce cytotoxicity.

### 2.14 Cell Transfection

A lentiviral transfection system was employed to knockdown or overexpress MXRA7 in the DC-2 cell line. Short hairpin RNAs targeting MXRA7 (shRNA^MXRA7^) and negative control short hairpin RNAs (shRNA^NC^) were designed, constructed, and integrated into the MISSION TRC2 pLKO.5-puro vector (Sigma-Aldrich). For overexpression experiments, the MXRA7-specific overexpression vector (EV-MXRA7) and the empty vector control (EV) were integrated into the pCDH-CMV-MCS-EF1α-Puro vector (SBI, USA). All the above vector construction procedures were performed by Wuhan Servicebio Technology Co., Ltd. (China).

Subsequently, lentiviruses were packaged to generate small interfering RNAs (siRNAs) and mediate exogenous MXRA7 expression. The sequence of siRNA^NC^ was TTCTCCGAACGTGTCACGTTT, and that of siRNA^MXRA7^was CTGAAGAAGCCGAGGAAGATT; the coding sequence of MXRA7 used in this study was verified to be correct by sequencing. DC-2 cells were separately infected with lentiviruses carrying shRNA^MXRA7^, shRNA^NC^, the MXRA7 overexpression vector, or the empty vector. The culture medium was replaced after overnight incubation, and the cells were then transferred to a selection medium containing 2 μg/mL puromycin (Sigma-Aldrich) for screening. The cells were maintained and passaged in the selection medium until subsequent experimental assays or cryopreservation, ensuring that stable MXRA7 knockdown or overexpression was achieved at all experimental time points.

### 2.15 RNA Extraction and qRT-PCR Analysis

Total RNA was isolated from murine epididymal tissues or DC-2 and PC-1 cells following the standard TRIzol extraction protocol. Using 1 μg of total RNA as the template, complementary DNA (cDNA) was synthesized with the PrimeScript RT Reagent Kit (containing gDNA Eraser, Takara Bio Inc., Japan) to eliminate genomic DNA contamination. Quantitative real-time polymerase chain reaction (qRT-PCR) assays were performed on the StepOnePlus Real-Time PCR System (Applied Biosystems, USA), with SYBR Green Premix Pro Taq HS (Accurate Biotechnology, China) used as the fluorescent dye.

Primers for MXRA7, IL-6, IL-1β, TNF-α, and the reference gene GAPDH were designed based on the corresponding gene sequences from the NCBI database and synthesized by Wuhan Servicebio (China). The 20 μL reaction system consisted of the following components: 10 μL SYBR Green Premix, 0.4 μL forward primer (10 μM), 0.4 μL reverse primer (10 μM), 2 μL cDNA template, and 7.2 μL nuclease-free water. The amplification conditions were set as follows: initial denaturation at 95 °C for 30 s, followed by 40 cycles of denaturation at 95 °C for 5 s and annealing/extension at 60 °C for 30 s. After amplification, a melting curve analysis (65–95 °C) was performed to verify the specificity of the PCR products. The relative mRNA expression levels of target genes were calculated using the 2^-ΔΔCt^ method and normalized against the reference gene GAPDH. The primer sequences were as follows: MXRA7 (NM_026280.3): Forward 5′-GGGCCACCCACTGAAGAAC-3′ , Reverse 5′-GTCTGACATCTCGCCAAAGGT-3′ ; IL-6 (NM_031168.2): Forward 5′-CAAGACTTCCATCAGTGC-3′ , Reverse 5′-GGCAGTCTCCTCATTGAATCC-3′; IL-1β (NM_008361.4): Forward 5′-TGGCAATGAGGATGACTTTCTTAC-3′ , Reverse 5′-GCTTGTGCTCTGCTGTGTGGT-3′; TNF-α (NM_013693.3): Forward 5′-CCTGTAGCCCACGTCTAGC-3′,Reverse 5′-GGTTTGCTACAACATGGGCTACA-3′.

### 2.16 Western blot (WB)

Epididymal tissues or cell pellets were homogenized by grinding in a lysis buffer containing RIPA buffer, PMSF protease inhibitor, and phosphatase inhibitors. After incubation on ice, the homogenates were centrifuged at 11,000 r·min⁻¹ for 15 min at 4 °C (centrifugal radius: 10 cm). The supernatant was collected to extract nuclear proteins and total tissue proteins.Protein concentrations were determined using the bicinchoninic acid (BCA) assay. Subsequently, the protein samples were boiled at 100 °C for 10 min, followed by sodium dodecyl sulfate-polyacrylamide gel electrophoresis (SDS-PAGE) procedures including gel preparation, sample loading, and electrophoresis. The separated proteins were transferred onto polyvinylidene fluoride (PVDF) membranes. The membranes were blocked with 5% non-fat milk on a shaking platform at room temperature for 1.5 h, then washed three times with TBST. Primary antibody working solutions were added, and the membranes were incubated overnight at 4 °C. The primary antibodies were recovered, and the membranes were washed three times with TBST again. HRP-conjugated secondary antibodies (1:10,000 dilution) were added and incubated for 1 h on a shaker. After three additional TBST washes, the membranes were treated with chemiluminescent substrate and exposed to X-ray film for visualization. The protein bands were quantified using Image J software.

### 2.17 Luciferase Reporter Assay

The firefly luciferase reporter plasmid pNFκB-luc and the reference plasmid pRL-TK (Beyotime Biotechnology, China) were co-transfected into cells using Lipofect5000 Prime DNA/siRNA Transfection Reagent (Biodai Biotechnology, China). At 48 h post-transfection, luciferase activity was measured in accordance with the manufacturer’s instructions for the Dual-Luciferase Reporter Assay Kit (Beyotime Biotechnology, China). The transcriptional activity of NF-κB was calculated as the ratio of firefly luciferase activity to Renilla luciferase activity(Liu et al., 2020).

### 2.18 Phos-tag SDS-PAGE

The extracted nuclear proteins and total tissue proteins were subjected to electrophoresis on precast Phos-PAGE gels (Abmart, China) to separate phosphorylated and non-phosphorylated proteins. After sample loading and electrophoresis, the gels were washed twice with methanol-free transfer buffer containing 5 mM ethylenediaminetetraacetic acid (EDTA) for 10 min per wash, followed by an additional wash with methanol-free transfer buffer alone. Subsequently, membrane transfer was performed, with all remaining procedures consistent with those of the standard WB assay.

### 2.19 Co-Immunoprecipitation (Co-IP)

Cells were lysed in ice-cold RIPA lysis buffer supplemented with protease and phosphatase inhibitors (Thermo Fisher Scientific) for 30 min. After centrifugation at 12,000 × g for 15 min at 4 °C, the supernatant was collected and incubated with 20 μL of protein A/G agarose beads (Santa Cruz Biotechnology) for 1 h at 4 °C with gentle shaking for pre-clearing. Subsequently, 2–5 μg of primary antibody or normal IgG (used as a negative control) was added to the pre-cleared lysate and incubated overnight at 4 °C, followed by the addition of 30 μL of protein A/G agarose beads and further incubation for 4 h at 4 °C. The beads were washed five times with cold PBS containing 0.1% Triton X-100. The immunoprecipitated complexes were resuspended in 2× SDS-PAGE loading buffer, and the target proteins were eluted by boiling for 10 min. Finally, the interacting proteins were detected via WB assay using specific antibodies.

### 2.20 In Vitro Phosphorylation Assay

The phosphorylation assay in vitro was performed as described previously(Xiao et al., 2024) with minor modifications. The MXRA7-containing complexes immunoprecipitated with MXRA7 antibody were washed twice with lysis buffer and once with wash buffer (40 mM HEPES pH 7.6, 10 mM MgCl₂, 1 mM EGTA, 1 mM DTT). The complexes were then incubated with recombinant active PKCα in kinase buffer (40 mM HEPES pH 7.6, 10 mM MgCl₂, 1 mM EGTA, 1 mM DTT, 2.5 mM β-glycerophosphate, 0.01% Triton X-100, 200 μM ATP) in the presence or absence of ATP at 30 °C for 30 min. The reaction was terminated by adding 5× Laemmli sample buffer, followed by heating at 95 °C for 5 min. Subsequently, the samples were subjected to Phos-tag SDS-PAGE and immunoblotted with MXRA7 antibody to detect the phosphorylation level of MXRA7.

### 2.21 Sperm In Vitro Incubation

The treated DC-2 cells were washed and incubated in serum- and antibiotic-free IMDM medium (without FBS and penicillin/streptomycin) for 24 h. The supernatant was collected by centrifugation at 1,000 r·min⁻¹ for 5 min. Spermatozoa were harvested from the cauda epididymidis of normal ICR mice and incubated at 37 °C for 30 min. The obtained samples were centrifuged at 400 × g for 5 min at room temperature to separate murine epididymal spermatozoa from epididymal fluid. The processed sperm samples were co-incubated with the collected cell supernatant for 12 h at room temperature under 5% CO₂ conditions, then processed and analyzed according to the aforementioned sperm treatment protocol.

### 2.22 Statistical Analysis

Data are presented as mean ± standard deviation (SD). Selected images were analyzed using ImageJ software. Statistical differences between two groups were determined by the Student’s t-test, while differences among multiple groups were assessed by one-way analysis of variance (ANOVA), followed by Tukey’s post-hoc test for multiple comparisons. A p-value < 0.05 was considered statistically significant.

## 3 Result

### 3.1 Exercise - Induced Fatigue Impairs Human Sperm Motility

To investigate the impact of EIF on male fertility, we measured sperm parameters in a cohort of EIF patients and compared the results with those of healthy controls (Fig. 1A). Serum biomarker analysis of EIF patients (Fig. 1B) revealed significant reductions in testosterone levels, metabolic waste clearance capacity, and antioxidant capacity. Assessment of sperm quality parameters (Fig. 1C) demonstrated that EIF patients exhibited marked declines in sperm motility-related traits, particularly in motility, straight-line velocity, and average path velocity. Immunofluorescence staining of sperm (Fig. 1D) was performed to evaluate acrosomal integrity (labeled by PNA), DNA damage (detected by TUNEL and 8-OHdG), and motility-related markers (α-tubulin, COX IV, and hexokinase I). The results indicated that EIF patients showed mild but non-significant acrosomal and DNA damage, whereas the expression of motility-associated markers was significantly downregulated. Collectively, these findings suggest that EIF exerts an adverse effect on sperm motility.

**Fig. 1.**
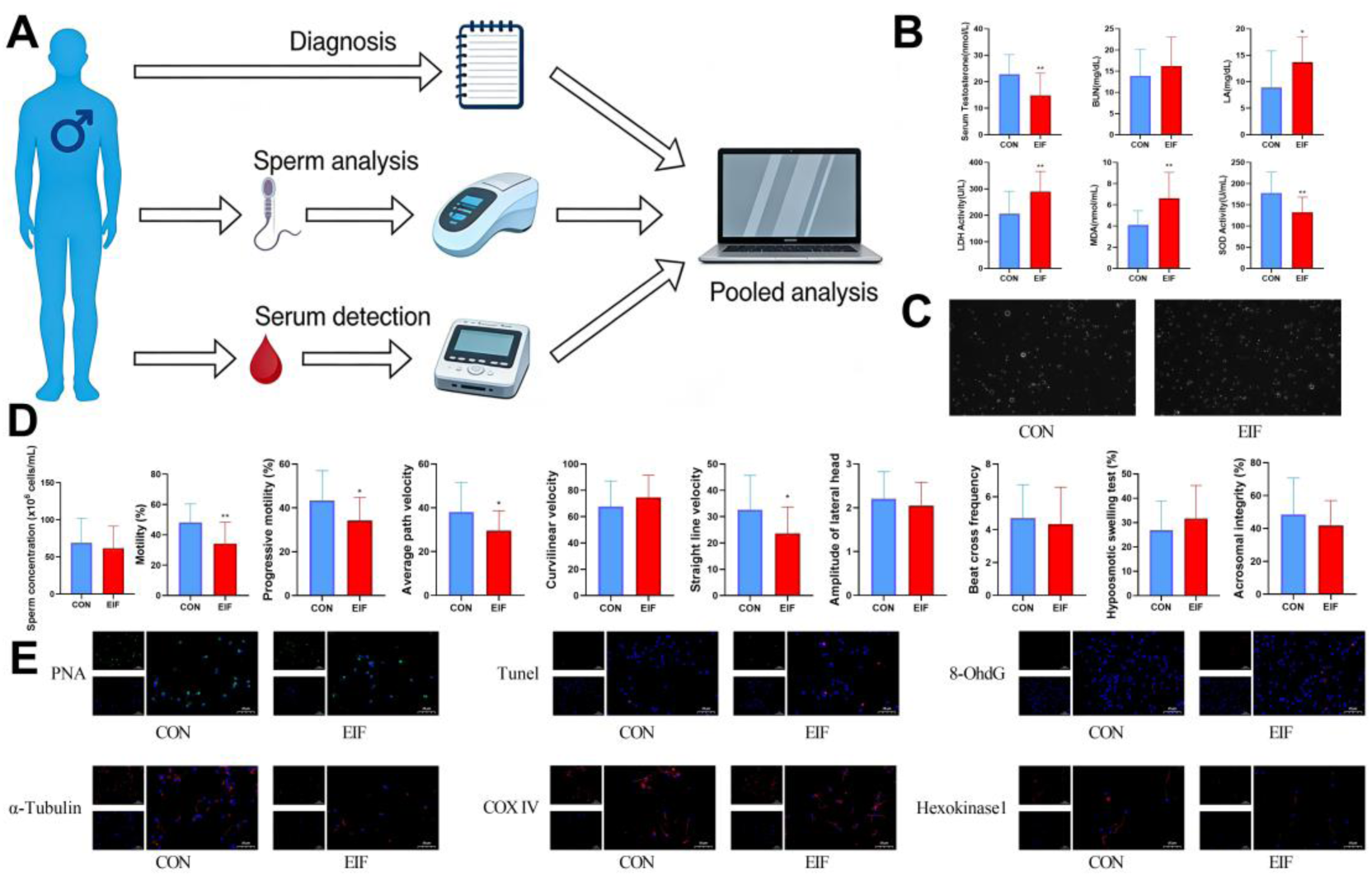
Sperm Motility of EIF Patients Is Compromised. Note: A: Research methods for human subjects; B: Serum detection levels in human subjects; C: Sperm quality parameters, acrosome integrity, and hypo-osmotic swelling test; D: Immunofluorescence staining of sperm smears: green fluorescence labels the acrosome, red fluorescence labels the target protein, and blue fluorescence indicates DAPI-stained cell nuclei; *p < 0.05, *p < 0.01.

### 3.2 Exercise - Induced Fatigue Impairs Mouse Sperm Quality

To further investigate the effects of EIF on sperm, we conducted animal experiments by establishing an EIF mouse model (Fig. 2A). Additionally, a vitamin C supplementation group was included to explore whether regular exercise and anti-fatigue supplements could exert beneficial effects on sperm quality. After the intervention period, EIF model mice exhibited significant decreases in body weight (Fig. 2B), hair luster and smoothness (Fig. 2C), and exercise capacity (Fig. 2D). Serum tests (Fig. 2E) assessing murine metabolic status (LA, BUN, LDH) and antioxidant levels (MDA and SOD) revealed that EIF mice had significantly increased metabolic waste accumulation and elevated oxidative stress levels. Histomorphological observations (Fig. 2F) showed obvious edema in the gastrocnemius muscle of EIF mice with increased myofiber spacing, indicating muscle fatigue. Collectively, these results confirmed the successful establishment of the EIF mouse model. Meanwhile, VC supplementation exerted a certain ameliorative effect, significantly alleviating the fatigue degree of EIF mice. Regarding the reproductive system of EIF mice (Fig. 2G-J), serum testosterone levels and epididymal index were significantly decreased. Changes in sperm quality were consistent with those observed in human EIF patients, primarily manifested as a significant decline in sperm motility. However, distinct from human findings, EIF mice also exhibited marked reductions in sperm concentration and acrosomal integrity. This discrepancy may be attributed to the fact that the forced swimming training in mice was closer to the physical exercise limit, resulting in a more severe fatigue degree compared to human subjects in this study. Immunofluorescence staining of mouse sperm smears showed a slight decrease in acrosomal integrity but no significant change in DNA damage levels (Fig. 2K). Similar to the human results, proteins associated with sperm motility displayed obvious impairment in EIF mice (Fig. 2L). Notably, vitamin C supplementation did not exert a significant protective effect on the reproductive system of EIF model mice. Although modest improvements were observed in the epididymal index and immunofluorescence intensity of sperm motility-related proteins, various sperm parameters indicated that vitamin C failed to significantly improve overall sperm quality. In summary, the above experiments confirmed that EIF impairs sperm quality, particularly sperm motility.

**Fig. 2.**
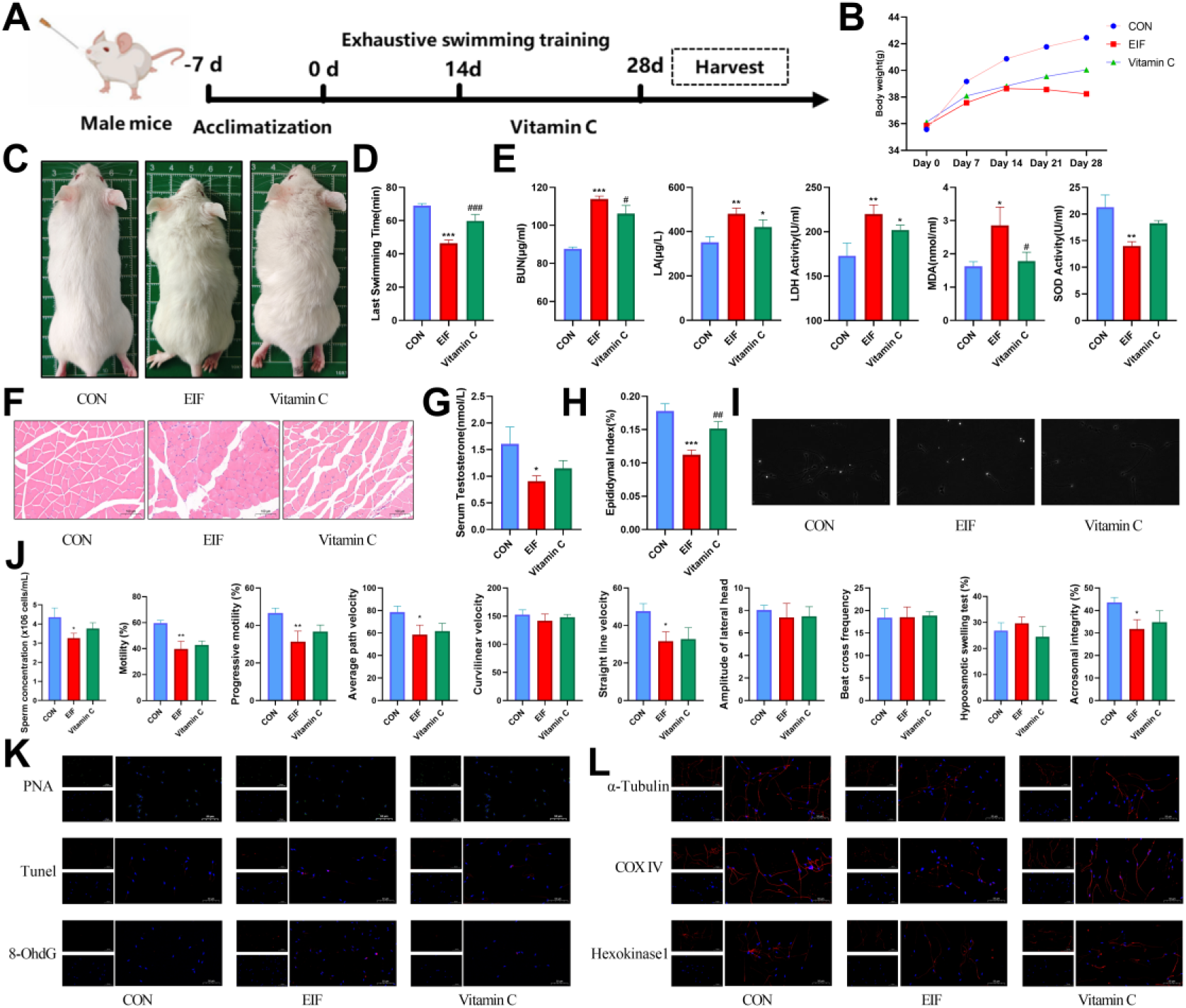
Sperm Quality of EIF Mice Is Significantly Impaired. Note: A: Animal experimental procedures, modeling and intervention time periods; B: Schematic diagram of body weight changes in mice; C: Morphological observation images of mice; D: Final swimming time of mice; E: Serum fatigue and antioxidant indicators in mice; F: Hematoxylin-eosin (HE) staining of gastrocnemius muscles; G: Serum testosterone levels in mice; H: Epididymal index (epididymal weight / total body weight); I: Sperm observation of mice; J: Sperm quality parameters, acrosome integrity and hypo-osmotic swelling test in mice; K: Immunofluorescence staining of sperm PNA and DNA damage-related markers: red fluorescence labels the target protein, and blue fluorescence indicates DAPI-stained cell nuclei; L: Immunofluorescence staining of proteins related to sperm motility; *P < 0.05, **P < 0.01, ***P < 0.001 vs. CON group; #P < 0.05, ##P < 0.01, ###P < 0.001 vs. EIF group.

### 3.3 Different Segments of the Epididymis in Exercise - Induced Fatigue Mice Exhibit Varying Degrees of Inflammation and Pyroptosis

Since the inflammatory level in the epididymal fluid of EIF mice was significantly elevated (Fig. 3A), we hypothesized that EIF might induce epididymitis and thus carried out studies focusing on three distinct regions of the epididymis: the caput, corpus, and cauda. Histological observations of the epididymis (Fig. 3B) showed that the epididymal walls in the caput, corpus, and cauda of EIF mice were all significantly thickened, with the most prominent alterations observed in the caput, indicating the occurrence of inflammation. Neutral α-glucosidase (NAG) is a functional epididymal marker that is almost exclusively derived from the epididymis and can reflect the integrity of epididymal function to a certain extent(Jin et al., 2024). Results of NAG immunohistochemistry (Fig. 3C) demonstrated impaired epididymal function in EIF mice. TUNEL staining of the epididymis (Fig. 3D) indicated obvious apoptosis in the caput and corpus epididymis of EIF mice. Immunofluorescence staining for IL-1β and qPCR detection of inflammatory factors (Fig. 3E–F) confirmed that a marked inflammatory response occurred in the epididymis of EIF mice. Further WB assays (Fig. 3G) confirmed that the epididymis of EIF mice not only underwent inflammation but also exhibited pyroptosis. Collectively, these experiments demonstrated that EIF induces epididymitis in mice, while vitamin C exerted a modest ameliorative effect on this pathological process. Through comprehensive analysis and literature retrieval, we proposed that MXRA7 might play a crucial role in EIF-induced epididymitis. Specifically, MXRA7 expression was significantly upregulated in the epididymis of EIF mice (Fig. 3F–G), and its expression levels varied remarkably across different epididymal segments (Fig. 3H–J). Multiple lines of evidence verified that the expression of MXRA7 in the cauda epididymis was significantly higher than that in the caput epididymis. Subsequent detailed detection of inflammatory and pyroptotic levels in distinct epididymal segments of EIF mice (Fig. 3K–L) revealed that the inflammatory level in the caput epididymis was higher than that in the corpus epididymis and significantly higher than that in the cauda epididymis. This expression pattern was inversely correlated with the segment-specific expression of MXRA7, implying a potential association between MXRA7 expression and epididymal inflammation.

**Fig. 3.**
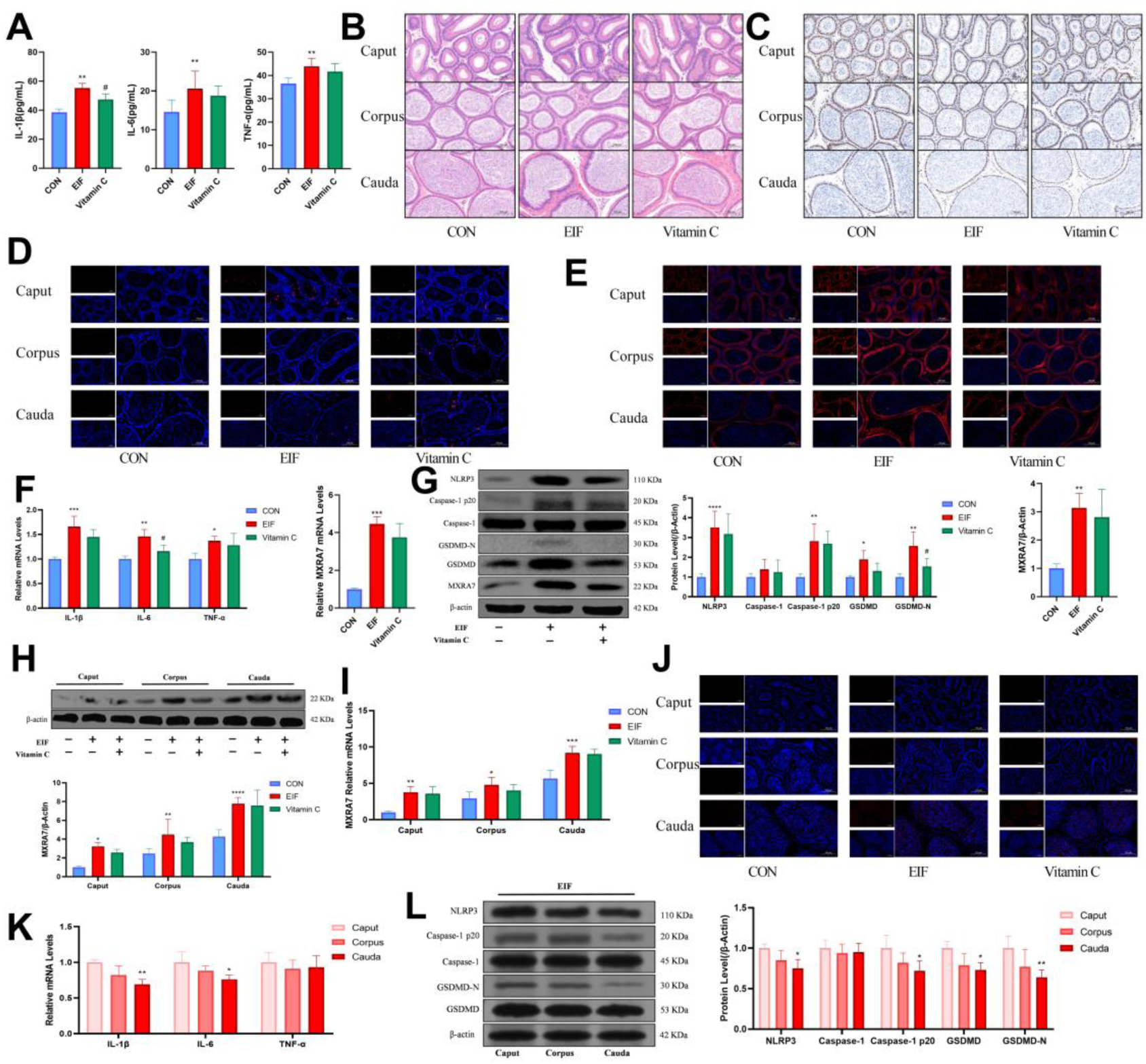
Different Segments of the Epididymis in EIF Mice Display Distinct Levels of Pyroptosis, Inflammation, and MXRA7 Expression. Note: A: Inflammatory cytokine levels in mouse epididymal fluid; B: Morphological observation of different epididymal regions in mice of each group by HE staining; C: Functional status observation of different epididymal regions in mice of each group by NAG immunohistochemical staining; D: Apoptosis observation of different epididymal regions in mice of each group by TUNEL staining; E: IL-1β immunofluorescence staining of different epididymal segments; F: mRNA expression levels of inflammatory cytokines and MXRA7 in mouse epididymis; G: Detection of pyroptosis-related proteins and MXRA7 protein expression levels in epididymis by WB; H: Detection of MXRA7 protein expression levels in different epididymal segments of mice in each group by WB; I: MXRA7 mRNA expression levels in different epididymal segments of mice in each group; J: MXRA7 immunofluorescence staining of different epididymal segments; K: mRNA expression levels of inflammatory cytokines in different epididymal segments of EIF mice; L: Detection of pyroptosis-related proteins in different epididymal segments of EIF mice by WB; *P < 0.05, **P < 0.01, ***P < 0.001, ****P < 0.0001 vs. CON group; #P < 0.05, ##P < 0.01, ###P < 0.001, ####P < 0.0001 vs. EIF group.

### 3.4 Significant Differences in LPS- and ATP-Induced Pyroptosis and MXRA7 Expression Between Caput and Cauda Epididymal Cells

To further verify the differences and potential correlation between epididymal caput and cauda cells in terms of pyroptosis and MXRA7 expression, as well as to exclude the interference of other factors such as the physical structural characteristics of animals, we cultured the PC-1 cell line derived from the epididymal caput and the DC-2 cell line derived from the epididymal cauda. Pyroptosis was induced in both cell lines using the same doses of ATP and LPS, and the pyroptosis profiles and MXRA7 expression differences between the two cell lines were observed. In terms of cell viability (Fig. 4A), the pyroptosis model of PC-1 cells exhibited a more statistically significant difference compared with CON group. With respect to inflammatory factors (Fig. 4B–C), the PC-1 cell line showed slightly more significant differences in IL-1β and IL-6 levels, whereas no obvious difference was detected in TNF-α levels. Immunofluorescence staining (Fig. 4D) showed that, compared with DC-2 cells, ATP-and LPS-induced PC-1 cells exhibited slightly higher fluorescence intensity in IL-1β and TUNEL staining, while the fluorescence intensity of NAG was slightly lower. No obvious change was observed in GSDMD immunofluorescence. Western blot (WB) analysis (Fig. 4E) revealed that DC-2 cells had lower levels of apoptosis and pyroptosis (as indicated by Caspase-1 p20 and GSDMD-N) compared with PC-1 cells, with statistically significant differences. Detection of MXRA7 levels (Fig. 4F–G) indicated that both the protein and mRNA levels of MXRA7 were significantly increased in DC-2 cells compared with PC-1 cells. Collectively, these results further confirmed that under the same doses of ATP and LPS, DC-2 cells exhibited lower degrees of apoptosis and inflammation than PC-1 cells, suggesting that the anti-inflammatory capacity of the epididymal cauda is stronger than that of the epididymal caput. Meanwhile, the segment-specific difference in MXRA7 expression was verified in more detail, and the association between MXRA7 and pyroptosis warrants further investigation.

**Fig. 4.**
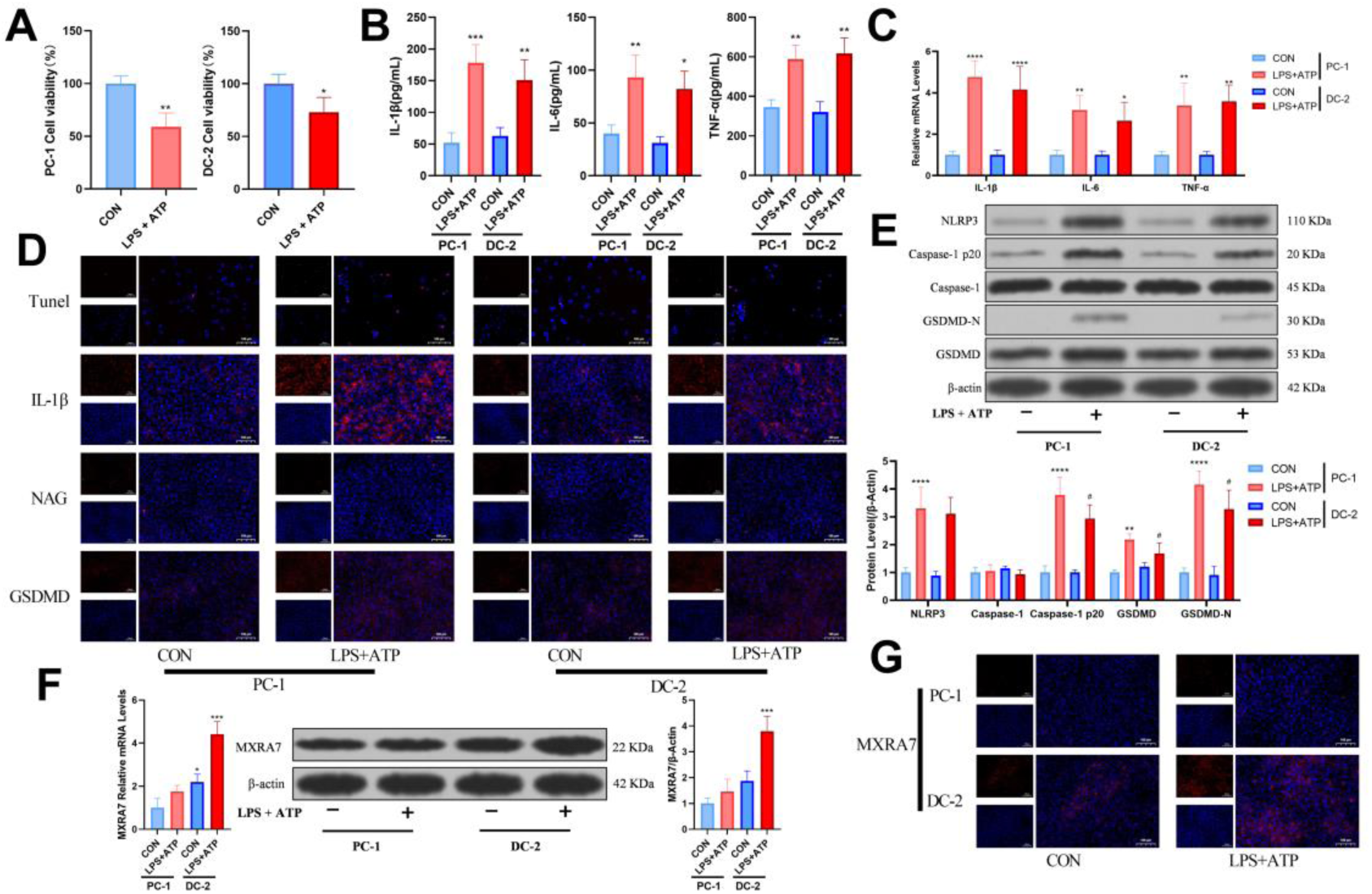
Differences Between PC-1 and DC-2 Cell Lines in Response to LPS and ATP. Note: A: Cell viability detected by CCK-8 assay; B-C: Inflammatory cytokine levels in cell supernatant and cellular mRNA levels; D: Immunofluorescence staining of PC-1 and DC-2 cells; E: Detection of pyroptosis-related proteins and quantitative indicators by WB; F-G: Detection of MXRA7 levels by WB, qPCR and immunofluorescence assay; *P < 0.05, **P < 0.01, ***P < 0.001, ****P < 0.0001 vs. the same cell line; #P < 0.05, ##P < 0.01, ###P < 0.001, ####P < 0.0001 vs. the same index in different cell lines.

### 3.5 Transcriptomics Analysis Reveals an Association Between MXRA7 and the NF-κB Signaling Pathway in Epididymitis

Given the potential association between MXRA7 and pyroptosis or inflammation, we performed transcriptomic analysis using epididymitis-related datasets (GSE145443, GSE199903) retrieved from the GEO database. Considering the data quality across multiple samples, cells with detected outliers or fewer than 500 expressed genes were filtered out, followed by the removal of doublets. A total of 2,611 cells were retained for subsequent analyses. Violin plots and scatter plots were generated using the filtered data (Fig. 5A–B), and 3,000 highly variable genes (HVGs) were identified (Fig. 5C).

**Fig. 5.**
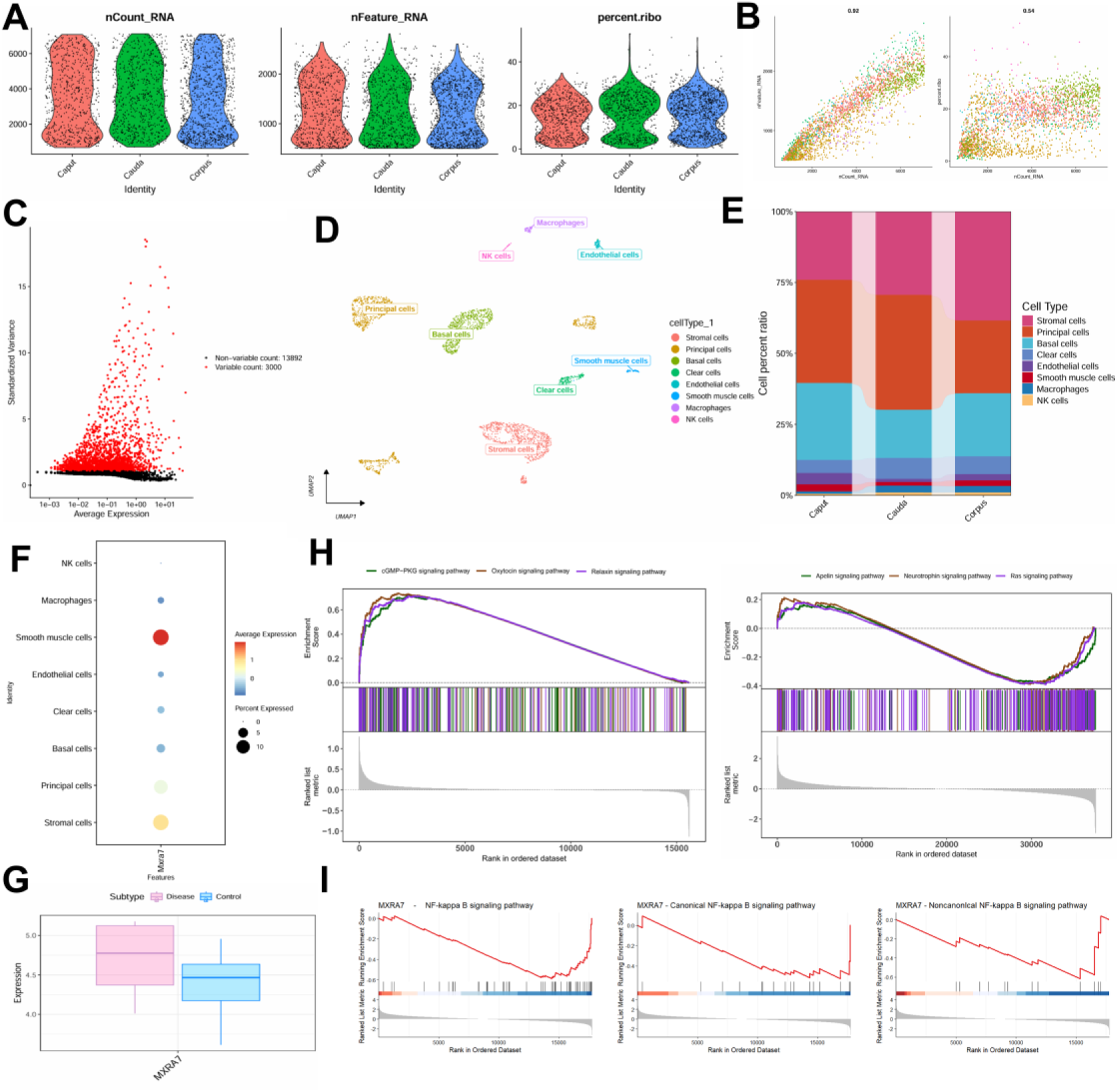
Transcriptomic Analysis of MXRA7 Expression in Epididymitis. Note: A-B: Quality control; C: 3,000 significantly upregulated genes; D: Eight annotated epididymal cell types identified by UMAP dimensionality reduction; E: Histogram of the proportions of different epididymal cell types across distinct epididymal segments; F: MXRA7 expression in various epididymal cell types; G: MXRA7 expression differences between normal epididymis and epididymitis samples; H: GSEA analysis of MXRA7 (left: GSE145443, right: GSE199903); I: GSEA analysis of MXRA7 on the NF-κB signaling pathway, as well as the canonical and non-canonical NF-κB signaling pathways.

Subsequently, the data were successively subjected to normalization, homogenization, PCA, and Harmony analysis. Following dimensionality reduction via UMAP, a total of 12 cell clusters were obtained. These clusters were annotated using known cell markers, and the 12 clusters were categorized into 8 cell types, namely Stromal cells, Principal cells, Basal cells, Clear cells, Endothelial cells, Smooth muscle cells, Macrophages, and NK cells (Fig. 5D). A bar chart was generated to visualize the proportions of these 8 cell types across different epididymal segments (Fig. 5E). Next, the Dotplot and FeaturePlot functions in the Seurat R package were used to examine the expression of MXRA7 at the single-cell level (Fig. 5F), and the differential expression of MXRA7 between normal and epididymitis samples was analyzed (Fig. 5G). The results showed that MXRA7 exhibited the highest expression in smooth muscle cells, principal cells, and clear cells within the epididymis. Given that epididymal epithelial cells are mainly composed of principal cells and clear cells, these findings further confirmed that the expression level of MXRA7 is increased in epididymitis. Subsequently, we performed gene set enrichment analysis (GSEA) on the GSE145443 and GSE199903 datasets (Fig. 5H). The results showed that MXRA7 was enriched in multiple signaling pathways, including the cGMP−PKG signaling pathway, Oxytocin signaling pathway, Relaxin signaling pathway, Apelin signaling pathway, Neurotrophin signaling pathway, and Ras signaling pathway. Through further analysis combined with existing literature(Sun et al., 2024), we hypothesized that MXRA7 might be associated with the NF-κB signaling pathway. A subsequent GSEA analysis focusing on the correlation between MXRA7 and the NF-κB signaling pathway confirmed a significant association between the two (Fig. 5I).

### 3.6 MXRA7 Inhibits the NF-κB Signaling Pathway and Attenuates Inflammation

To investigate the specific role of MXRA7, we knocked down or overexpressed MXRA7 in the DC-2 cell line using a lentiviral infection system. The MXRA7-knockdown and MXRA7-overexpression models of DC-2 cells were successfully established and validated (Fig. 6A–B). Interestingly, the results of cell viability assays (Fig. 6C) showed that MXRA7 knockdown severely impaired cell proliferation, with an effect even stronger than that observed in the pyroptosis model, whereas MXRA7 overexpression significantly enhanced cell viability, with an effect superior to that of the CON group. These findings indicate that MXRA7 plays a crucial role in cell proliferation and differentiation, which is consistent with the conclusions of previous studies(Ma et al., 2023, Sun et al., 2023). Supernatants and cellular RNA were collected to determine the levels of inflammatory factors (Fig. 6D–E). The results showed that MXRA7 knockdown promoted inflammation, whereas MXRA7 overexpression inhibited ATP- and LPS-induced inflammation. Meanwhile, MXRA7 knockdown also induced cell apoptosis and increased IL-1β levels, while MXRA7 overexpression exerted an inhibitory effect on these processes; however, no significant effect on NAG was observed (Fig. 6F–H). Detection of the NF-κB signaling pathway (Fig. 6I–J) revealed that MXRA7 inhibited the NF-κB signaling pathway, exerting certain effects on p65 activation and nuclear translocation as well as p52/p100 expression. Notably, the inhibitory effect on p52/p100 was highly significant, indicating that MXRA7 primarily represses the non-canonical NF-κB pathway. Subsequent pNFκB-luc luciferase assays (Fig. 6K) further confirmed that MXRA7 can inhibit the NF-κB signaling pathway.

**Fig. 6.**
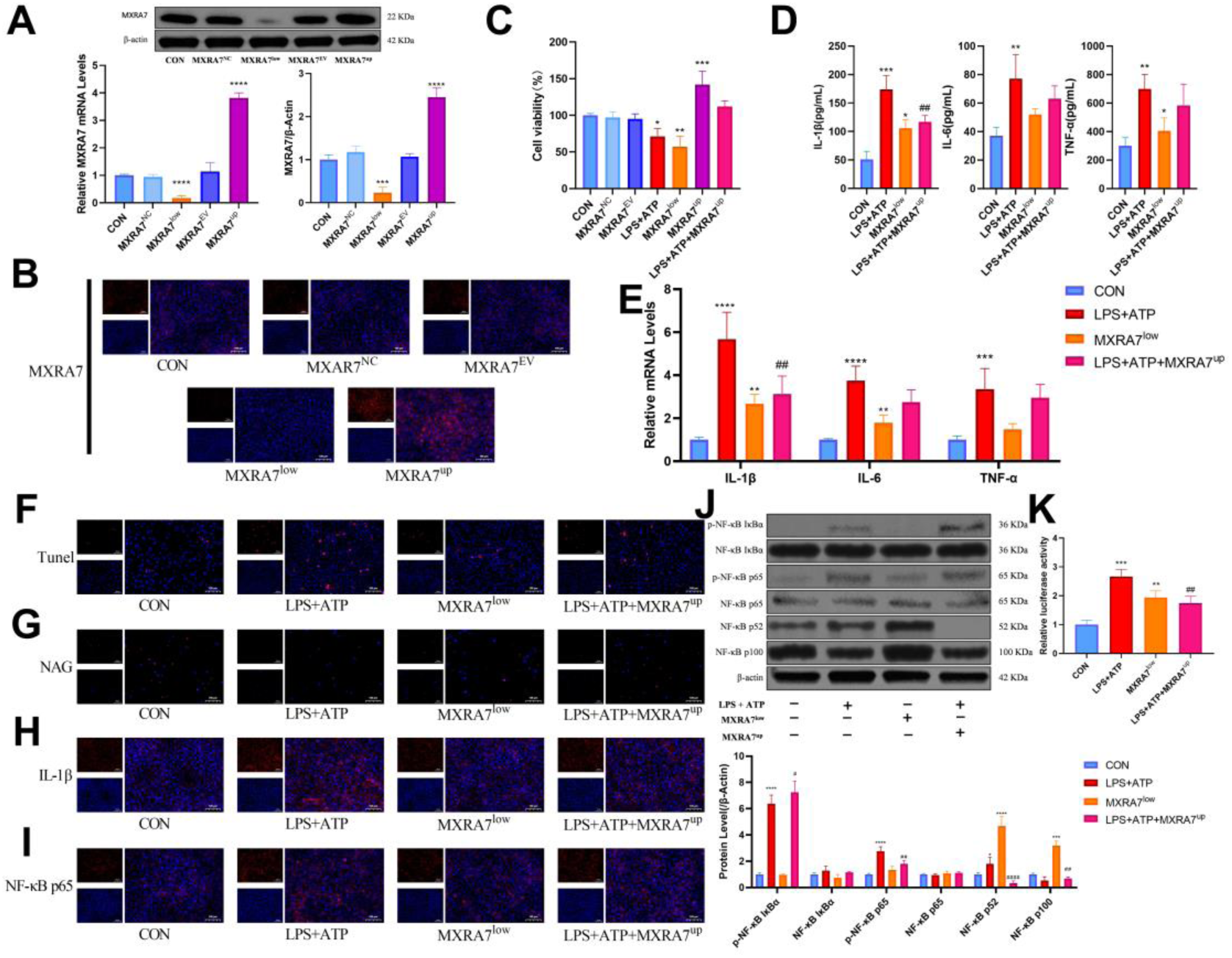
MXRA7 Inhibits the NF-κB Signaling Pathway Predominantly Mediated by the Non-Canonical Pathway. Note: A: MXRA7 protein expression and mRNA levels; B: MXRA7 immunofluorescence staining of DC-2 cells; C: DC-2 cell viability detected by CCK-8 assay; D-E: Inflammatory cytokine levels in cell supernatant and cellular inflammatory cytokine mRNA levels; F-I: Immunofluorescence staining of DC-2 cells; J: Detection of key proteins in the NF-κB signaling pathway by WB; K: pNFκB-luc luciferase reporter assay, with results expressed as the ratio of firefly luciferase activity to Renilla luciferase activity; *P < 0.05, **P < 0.01, ***P < 0.001, ****P < 0.0001 vs. CON group; #P < 0.05, ##P < 0.01, ###P < 0.001, ####P < 0.0001 vs. LPS+ATP group.

### 3.7 MXRA7 Is Phosphorylated by Activated PKCα

Through further analysis combined with recent studies on MXRA7, we hypothesized that MXRA7 might be associated with PKCα. Phorbol 12-myristate 13-acetate (PMA) is a PKCα agonist that directly activates PKCα via drug-mimicking effects with high specificity; BIM 1 is a specific inhibitor of PKCα. On the basis of pyroptosis induced by LPS + ATP, BIM 1 was added for intervention, with PMA intervention set as the control (Fig. 7A). The results showed that MXRA7 levels were significantly increased under PMA treatment, whereas BIM 1 exerted a marked inhibitory effect on MXRA7. These findings indicate that PKCα can enhance the expression level of MXRA7. Given that PKCα is a phosphorylation kinase, apart from the increase in expression level, MXRA7 may also undergo phosphorylation under the regulation of activated PKCα, which thus warrants further investigation for confirmation. MXRA7 phosphorylation and non-phosphorylation forms were separated via Phos-tag SDS-PAGE using DC-2 cell lysates (Fig. 7B). The results confirmed that MXRA7 is indeed phosphorylated, but only under LPS + ATP or PMA treatment conditions, suggesting that MXRA7 phosphorylation is associated with PKCα activation. To further verify the interaction between PKC and MXRA7, CO-IP assays were performed (Fig. 7C). The results showed that the interaction between phosphorylated PKCα (p-PKCα) and MXRA7 was the most pronounced. Subsequently, in vitro phosphorylation validation of MXRA7 by activated PKCα was conducted, followed by Phos-tag SDS-PAGE electrophoresis (Fig. 7D). These results once again confirmed that activated PKCα induces MXRA7 phosphorylation. On this basis, detection of the NF-κB signaling pathway was carried out (Fig. 7E–G) to verify the effect of PKCα on the NF-κB signaling pathway in the DC-2 cell line. It was found that PKCα appeared to exert a certain inhibitory effect on p52/p100, which might be related to phosphorylated MXRA7.

**Fig. 7.**
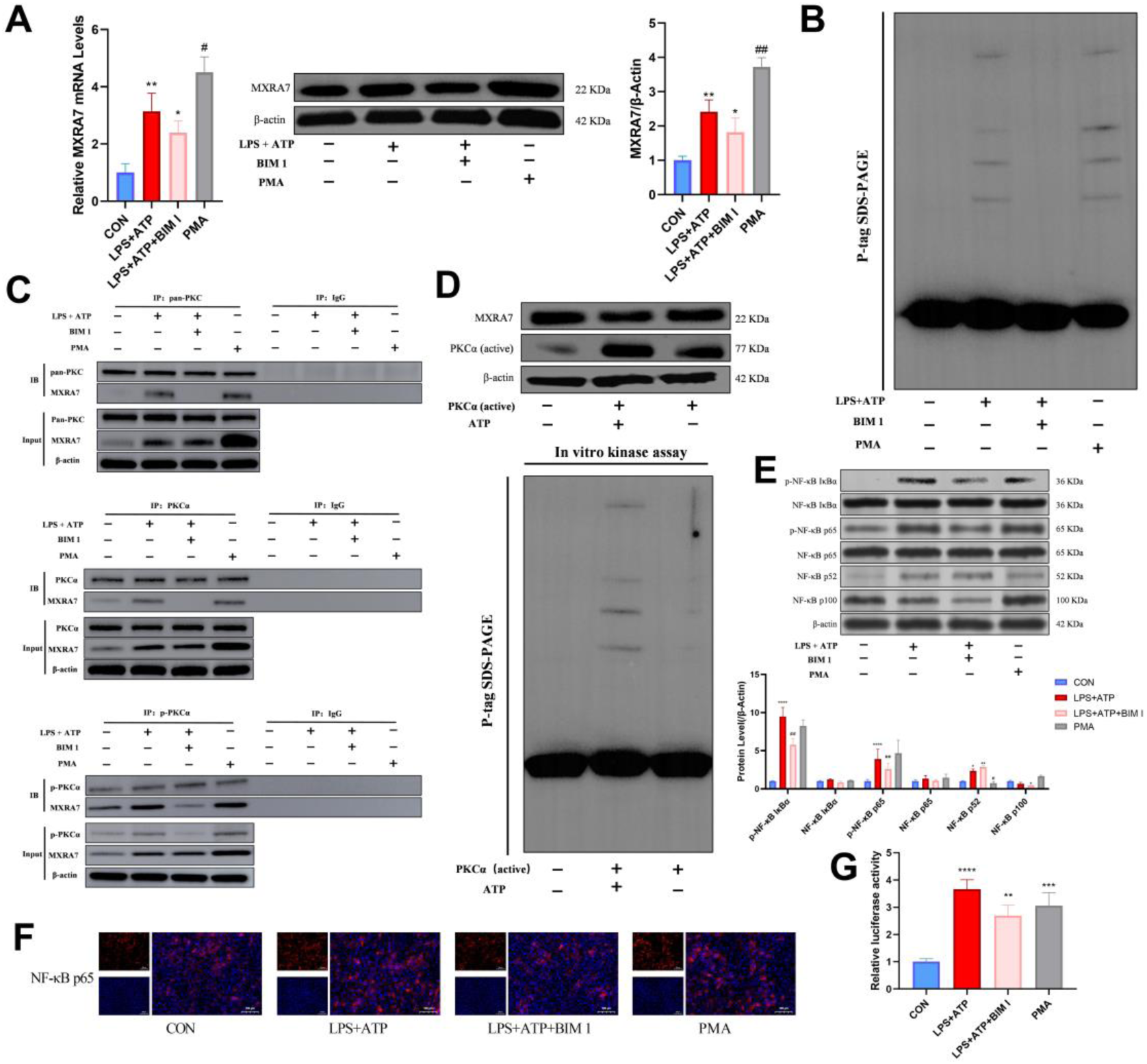
Activated PKCα Phosphorylates MXRA7. Note: A: MXRA7 protein expression and mRNA levels; B: Separation of phosphorylated and non-phosphorylated MXRA7 proteins by Phos-tag SDS-PAGE electrophoresis; C: Detection of the interaction between pan-PKC, PKCα, p-PKCα and MXRA7 by CO-IP assay; D: In vitro phosphorylation assay of MXRA7 with activated PKCα, and verification of MXRA7 phosphorylation by Phos-tag SDS-PAGE; E-F: Detection of key proteins in the NF-κB signaling pathway by WB and immunofluorescence staining; G: pNFκB-luc luciferase reporter assay, with results expressed as the ratio of firefly luciferase activity to Renilla luciferase activity; *P < 0.05, **P < 0.01, ***P < 0.001, ****P < 0.0001 vs. CON group; #P < 0.05, ##P < 0.01, ###P < 0.001, ####P < 0.0001 vs. LPS+ATP group.

### 3.8 MXRA7 Attenuates the Adverse Effects of Epididymal Epithelial Cell Pyroptosis on Sperm Motility

Although the aforementioned experiments have verified the role of MXRA7 in epididymal epithelial cell inflammation, considering that MXRA7 is a secreted protein, its effect on sperm under the condition of epididymitis remains to be further validated. DC-2 cells with MXRA7 knockdown or overexpression were induced to undergo pyroptosis, thoroughly washed, and their supernatants were collected and incubated with sperm from normal ICR mice to observe sperm status (Fig. 8A). After incubation, the supernatants were collected to detect inflammatory factors (Fig. 8B). The results showed that the levels of inflammatory factors were inhibited by MXRA7 to a certain extent. Post-incubation sperm were analyzed using an automatic sperm analyzer (Fig. 8C–D). Sperm incubated with supernatants from pyroptosis-induced DC-2 cells exhibited a similar phenotype to that observed in EIF mouse sperm in the aforementioned experiments, with sperm motility significantly reduced. In contrast, MXRA7 overexpression markedly alleviated the decline in sperm motility, whereas MXRA7 knockdown led to a significant inhibition of sperm motility. Results of immunofluorescence staining (Fig. 8E–J) further demonstrated that the regulatory effect of MXRA7 was mainly reflected in the improvement of sperm motility-related parameters. Interestingly, MXRA7 also appeared to exert a protective effect on the acrosome. In conclusion, MXRA7 exerts a definite protective effect on sperm, particularly on sperm motility, which may be associated with the function of epididymal epithelial cells in maintaining sperm viability.

**Fig. 8.**
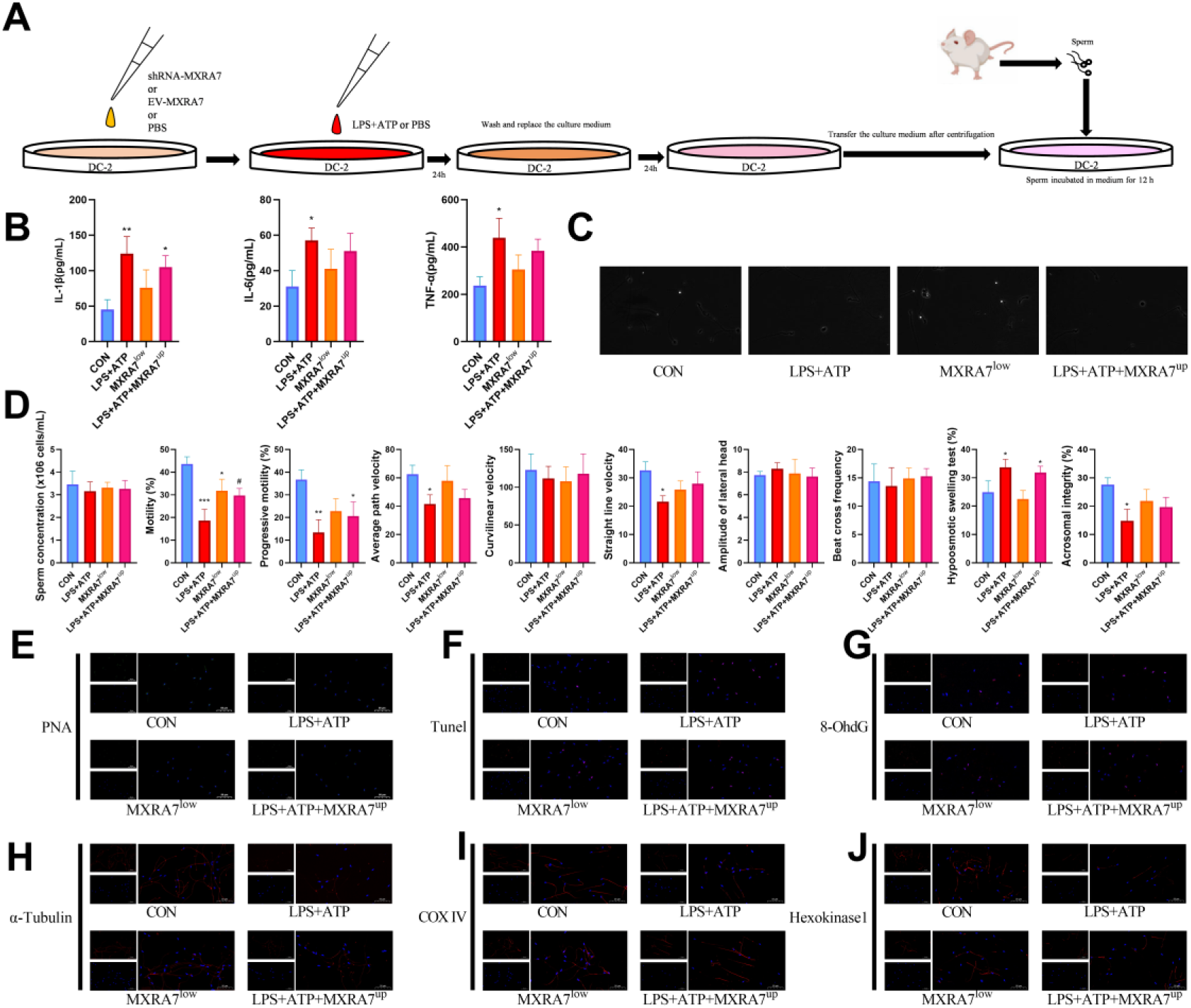
MXRA7 Ameliorates the Adverse Effects of Epididymal Epithelial Cell Pyroptosis on Sperm. Note: A: Schematic diagram of sperm in vitro incubation; B: Inflammatory cytokine levels in the supernatant after incubation; C-D: Analysis of sperm quality by automatic sperm analyzer; E-J: Detection of acrosome, DNA damage and motility-related indicators of sperm after incubation by immunofluorescence staining; *P < 0.05, **P < 0.01, ***P < 0.001, ****P < 0.0001 vs. CON group; #P < 0.05, ##P < 0.01, ###P < 0.001, ####P < 0.0001 vs. LPS+ATP group.

## 4 Discussion

Moderate exercise can promote human health, enhance physical fitness, prevent and treat various chronic diseases, improve mental state and elevate quality of life. However, chronic EIF caused by excessive exercise is not only detrimental to physical fitness improvement, but also induces adverse reactions in multiple organ systems of the body(Festa et al., 2023). To date, numerous studies have been conducted on EIF(Li et al., 2025, Kim et al., 2025, Xu et al., 2025, Yang et al., 2024a), yet most of them have focused on the musculoskeletal and nervous systems. Obviously, as a systemic and multisystem response, EIF is likely to exert impacts on other various systems, among which its effects on reproduction are also multifaceted(Tang et al., 2025). In the present study, we focused on the association between EIF and the reproductive system. To verify the adverse effects of EIF on sperm, we recruited volunteers with EIF symptoms, collected their semen and serum samples, and compared these samples with those from healthy volunteers. The results showed that sperm motility was indeed impaired and the inflammatory level of seminal plasma was significantly increased. This clinical study directly and practically confirmed that EIF impairs the reproductive system, laying a foundation for subsequent experiments.

Based on the confirmation from clinical experiments, we conducted animal experiments for further in-depth research. First, we induced EIF symptoms in mice, followed by sperm and serum tests similar to those in clinical studies. The results also showed that the sperm motility of EIF mice was significantly impaired. However, unlike the clinical findings, the sperm concentration and acrosome integrity of EIF mice were also significantly decreased. This discrepancy may be attributed to the fact that the EIF modeling in animal experiments resulted in a more severe degree of fatigue than that in clinical volunteers. In addition, we added vitamin C as an intervention supplement for EIF in animal experiments. Planned vitamin C supplementation during exercise is commonly used as a strategy to prevent and treat EIF(Rogers et al., 2023) and is a frequently employed supplement in EIF modeling-related studies(Chen et al., 2022). Nevertheless, the results indicated that although vitamin C alleviated EIF-induced fatigue, its ameliorative effect on sperm quality was relatively weak, suggesting that more targeted drugs may be required.

Unlike human volunteers, sperm samples from mice were collected directly from the epididymis. However, epididymal sperm also exhibited impairment, which might be attributed to the inflammatory factors released by epididymitis that affected epididymal sperm function. Subsequent in-depth studies on the epididymis revealed that the epididymis of EIF mice developed inflammation. Neutral α-glucosidase (NAG) can reflect the capacity of the epididymis to maintain sperm normalcy to a certain extent(Jin et al., 2024), and its level in the epididymis of EIF mice was also decreased. Further analysis was performed to examine whether pyroptosis occurred in the epididymis of EIF mice. NLRP3, Caspase-1, and GSDMD are core molecules of cellular pyroptosis, which function in the sequential order of “activation-transduction-execution”: as an inflammasome sensor, NLRP3 assembles and activates upon stimulation by pathogens, damage signals, and other stimuli; activated NLRP3 recruits and activates downstream Caspase-1; activated Caspase-1 cleaves the GSDMD protein, enabling its N-terminal domain to insert into the cell membrane and form pores, ultimately leading to cell rupture and death, namely pyroptosis (Wang et al., 2024, Broz, 2025). The results confirmed that the epididymis of EIF mice indeed developed inflammation and underwent pyroptosis.

Owing to the differences in cellular composition across distinct segments, the epididymis can be generally divided into three regions: the caput, corpus, and cauda(Sullivan et al., 2019, James et al., 2020). We performed separate detections on these three segments of the epididymis in EIF mice and found that the levels of apoptosis and inflammation in the caput epididymis appeared to be higher than those in the corpus and cauda. Further detection revealed that the degree of pyroptosis in the caput epididymis was significantly higher than that in the cauda. Given that the caput epididymis has a more complex morphological structure, physical damage such as friction caused by exercise may be more severe in this region. Therefore, we cultured mouse epididymal caput epithelial cells (PC-1) and mouse epididymal cauda epithelial cells (DC-2) in vitro, and induced pyroptosis by adding LPS and ATP(Yang et al., 2024b). The results still showed that the levels of pyroptosis and inflammation in DC-2 cells were lower than those in PC-1 cells, suggesting that this difference was not merely caused by variations in physical structure.

MXRA7 is an autocrine factor, and recent studies have indicated its association with immunity, inflammation, and cellular repair(Sun et al., 2023, Shen et al., 2023). We serendipitously discovered that the expression level of MXRA7 varies across the caput, corpus, and cauda of the epididymis, with remarkably higher expression in the cauda than in the caput; additionally, MXRA7 expression is upregulated in the epididymis of EIF mice. Further studies revealed that MXRA7 expression also exhibits differential patterns in cultured cells in vitro and is significantly increased upon stimulation with LPS and ATP. MXRA7 is presumably associated with epididymitis, and the relatively low inflammatory level in the epididymal cauda may be related to MXRA7. Transcriptomic analysis using public databases identified a significant association between MXRA7 and the NF-κB signaling pathway, suggesting that MXRA7 is likely to regulate inflammation and pyroptosis via the NF-κB signaling pathway(Afonina et al., 2017). Subsequent knockdown and overexpression experiments of MXRA7 in the DC-2 cell line confirmed that MXRA7 exerts an inhibitory effect on the NF-κB signaling pathway (predominantly the non-canonical pathway) as well as on inflammation.

Protein kinase C (PKC) is a family of calcium ion- and phospholipid-dependent protein kinases that are widely involved in the regulation of cell proliferation, differentiation, apoptosis, inflammatory responses, and signal transduction(Gada and Logothetis, 2022). The PKC family consists of multiple isoforms, among which PKCα belongs to the conventional PKC subtype. PKCα can regulate NLRP3 inflammasome activation, cellular pyroptosis progression, and epithelial cell function(Fanourakis et al., 2023, Shio et al., 2015, Zhou et al., 2024); it is directly associated with the NF-κB signaling pathway and can activate the canonical NF-κB signaling pathway(Wang et al., 2025). Based on an analysis of recent studies(Sun et al., 2023), we hypothesized that MXRA7 has a strong association with PKC and exhibits characteristics consistent with those of downstream molecules of PKC. MXRA7 is not only directly regulated by PMA (a PKCα agonist(Zhou et al., 2024)) and BIM 1 (a PKCα inhibitor(Suryawanshi et al., 2022)), but also exists in a phosphorylated state. In vitro kinase assays and co-immunoprecipitation (CO-IP) experiments between MXRA7 and PKC confirmed that activated PKCα exerts a phosphorylating effect on MXRA7. However, it must be acknowledged that since PKC can also affect the expression level of MXRA7, the specific mechanism underlying the action of phosphorylated MXRA7 requires further investigation. In conclusion, MXRA7 primarily inhibits the non-canonical NF-κB signaling pathway. Therefore, under external stimuli such as inflammation, activated PKCα activates the canonical NF-κB signaling pathway while simultaneously inhibiting the non-canonical NF-κB signaling pathway via MXRA7. MXRA7 also exerts a certain inhibitory effect on the canonical pathway, which may represent a negative feedback mechanism. The presence of MXRA7 reduces the NLRP3 transcription induced by activated NF-κB, thereby attenuating pyroptosis. Finally, co-incubation of epididymal sperm from normal mice with the DC-2 cell line demonstrated the direct effect of MXRA7 on sperm. Evidently, MXRA7 can effectively alleviate the adverse effects of DC-2 cell pyroptosis on sperm, and also verified in vitro that epididymitis and pyroptosis occurring in EIF mice cause damage to sperm. However, MXRA7 is also a secreted factor, and further in-depth research is still needed to clarify whether MXRA7 secreted by epididymal cauda epithelial cells can directly maintain sperm motility.

## 5 Conclusion

In summary, EIF induces pyroptosis in the epididymis, triggers epididymitis, and ultimately impairs sperm motility. Among the three segments of the epididymis (caput, corpus, and cauda), there are distinct differences in the degrees of inflammation and pyroptosis, with the lowest inflammatory level observed in the cauda. This regional difference is presumably associated with the high expression level of MXRA7 in the epididymal cauda. MXRA7 can alleviate epididymitis by inhibiting the NF-κB signaling pathway, and meanwhile, MXRA7 undergoes phosphorylation under the regulation of activated PKCα. During the process of EIF-induced epididymitis, MXRA7 is likely to exert a protective role, reducing the degree of pyroptosis mainly in epididymal cauda epithelial cells and thereby mitigating the damage to sperm quality to a certain extent.

## Conflict of interest statement

### Ethics approval

Regarding studies involving human subjects: This study was approved by the Institutional Ethics Committee (Approval No.: HBZY2023-C10-02). All experiments were performed in accordance with the principles of the Declaration of Helsinki of the World Medical Association, and informed consent as well as voluntary cooperation were obtained from each participant; Regarding animal experiments: All experimental procedures were strictly performed in accordance with the Guidelines for the Care and Use of Laboratory Animals and were approved by the Institutional Animal Care and Use Committee of Hubei University of Chinese Medicine (Approval No.: HUCMS55714520).

## Authors’ contributions

KT authored the article; XJ revised and reviewed it; ZF and XY provided platform support; XH and JL conducted validation and provided suggestions; MZ and YL performed the experimental reproducibility verification and cross-validation of data results; JC, YZ and MX provided the ideas and concepts.

